# Antimetabolic cooperativity with the clinically approved kidrolase and tyrosine kinase inhibitors to eradicate cml stem cells

**DOI:** 10.1101/2020.09.21.305714

**Authors:** Anne Trinh, Raeeka Khamari, Quentin Fovez, François-Xavier Mahon, Béatrice Turcq, Didier Bouscary, Patrice Maboudou, Marie Joncquel, Valérie Coiteux, Nicolas Germain, William Laine, Salim Dekiouk, Bart Ghesquiere, Thierry Idziorek, Bruno Quesnel, Jerome Kluza, Philippe Marchetti

**Affiliations:** Univ. Lille, CNRS, Inserm, CHU Lille, Institut de Recherche contre le Cancer de Lille, UMR9020 – UMR-S 1277 - Canther – Cancer Heterogeneity, Plasticity and Resistance to Therapies, F-59000 Lille, France; Institut Bergonié, Université de Bordeaux, CNRS SNC5010, Inserm, U1218 ACTION F – 33076 Bordeaux, France; Université de Paris, Institut Cochin, CNRS UMR8104, INSERM U1016, Paris, France; Centre de Bio-Pathologie, Banque de Tissus, CHU Lille, France; Assistance Publique-Hôpitaux de Paris. Centre-Université de Paris, Service d’Hématologie clinique, Hôpital Cochin, Paris, France; Department of Oncology and VIB, KU Leuven, Leuven, Belgium

**Keywords:** synthetic lethality, metabolic addiction, LSC, metabolic stress, stem-like cells

## Abstract

Long-term treatment with tyrosine kinase inhibitors (TKI) represents an effective treatment for chronic myeloid leukemia (CML) and discontinuation of TKI therapy is now proposed to patient with deep molecular responses. However, evidence demonstrating that TKI are unable to fully eradicate dormant leukemic stem cells indicate that new therapeutic strategies are needed to prevent molecular relapses. We investigated the metabolic pathways responsible for CML surviving to Imatinib exposure and its potential therapeutic utility to improve the efficiency of TKI against CML stem cells. Using complementary cell-based techniques, we demonstrated that TKI suppressed glycolysis in a large panel of BCR-ABL1 + cell lines as well as in primary CD34+ stem-like cells from CML patients. However, compensatory glutamine-dependent mitochondrial oxidation supported ATP synthesis and CML cell survival. Glutamine metabolism was inhibited by L-asparaginases such as Kidrolase without inducing predominant CML cell death. Clinically relevant concentrations of TKI render CML progenitors and stem cells susceptible to Kidrolase. The combination of TKI with L-asparaginase reactivated the intinsic apoptotic pathway leading to efficient CML cell death. Thus, targeting glutamine metabolism with the clinically-approved drug Kidrolase, in combination with TKI that suppress glycolysis represents an effective and widely applicable therapeutic strategy for eradicating CML stem cells.

## Introduction

With the development of tyrosine kinase inhibitor (TKI) therapy, the outcome of chronic myeloid leukemia (CML) patients has changed drastically. Unfortunately, TKI, such as Imatinib mesylate and other second or third generation of BCR-ABL1 inhibitors, target preferentially differentiated cells and leave some of the CML stem cells alive[1,2]. Indeed, a fraction of LSC can survive independently of BCR-ABL1 signaling and thus are totally insensitive to Imatinib [1]. TKIs are effective in inducing a long-term response and discontinuation of treatment is proposed to patients with a persistent deep molecular response [3,4]. The STIM study has shown that 40% of patients successfully achieved treatment free remission with no recurrence of the disease. However, resumption of treatment is necessary for patients who exhibit BCR-ABL1 transcript increase following Imatinib discontinuation. The inability of TKI to kill LSCs and/or progenitor cells is at the origin of relapses [5]. Thus, strategies to improve the efficiency of TKI against LSC in CML are needed to definitely eradicate the disease and allow long definitive TKI discontinuation in most patients.

Cancer cell metabolism has opened up a new avenue in cancer treatment because it appears possible to target specific metabolic features of cancer cells, thus providing a potential therapeutic window. Indeed, many oncogenes and oncogenic pathways that drive cancer development also drive metabolism. Classically, it is assumed that cancer metabolism is dependent on glycolysis that is predominant even in normoxic conditions (a phenomenon also called the Warburg effect). Thus, the aberrant activation of the MAPK pathway via the BRAFV600E mutation increases glucose uptake and glycolysis allowing intense cell proliferation (for review [6,7]). Consistent activation of the PI3K/Akt/mTOR pathway by BCR-ABL1 also increases glucose metabolism in leukemic cells [8,9]. Given these conditions, it is not surprising that the exposure to oncogenic “driver” inhibitors such as BRAFV600E inhibitors [10] or BCR-ABL1 inhibitors [11] dramatically reduces glucose uptake and glycolysis to promote cell cycle arrest.

However, it has become evident that cancer metabolism cannot be resumed to the Warburg effect and represents a more complex network linking glucose metabolism and others nutrients such as glutamine. Thus, numerous evidence indicate that mitochondrial oxidative metabolic pathways have a crucial role in cancer development particularly to immediately compensate for glucose deprivation. Interestingly, the dependence on mitochondrial oxidative metabolism allows cells to avoid cell death induced by MAPK inhibitors [7] or TKI [12]. Moreover, CML stem cells are particularly sensitive to mitochondrial oxidative metabolism inhibitors [13]. Thus, one interesting therapeutic strategy in cancer could be the combination of molecular-targeted drugs with antiglycolytic activities and inhibitors of the compensatory mitochondrial oxidative pathways thereby creating an “antimetabolic cooperativity” [6]. Herein, our objective was to develop a pre-clinical proof of concept of antimetabolic cooperativity against CML stem cells. We demonstrated that the combination of Kidrolase, a L-asparaginase used as treatment of acute lymphoblastic leukemia, with Imatinib possesses antimetabolic cooperativity and acts synergistically to eradicate CML stem cells *in vitro* and *ex vivo*.

## Materials and methods

### Chemicals

Imatinib mesylate was purchased from Sigma-Aldrich, Nilotinib and Dasatinib were purchased from Selleckchem. Kidrolase and Erwinase were provided by CHU Lille, [^13^C_6_] Glucose and [^13^C_5_] Glutamine were obtained from Professor Bart Ghesquière.

### Patients samples

CML samples were blood or bone marrow samples obtained from individuals in chronic or acute phase CML recruited from the Department of Haematology (Lille CHU, France), with informed consent in accordance with the Declaration of Helsinki and approval of the institutional ethical committee (CPP Lille). CD34+ cells were isolated from umbilical cord blood using the EasySep^™^ Human CD34 Positive Selection Kit II (Stemcell Technologies). Cytapheresis of CML patients were kindly provided by Pr. François-Xavier Mahon (CHU Bordeaux, France). CD34+ cells were cultivated at 37°C and 5% CO_2_ in RPMI medium (Gibco) supplemented with 10% fetal calf serum (Gibco), 50 U/ml penicillin, 50 mg/ml streptomycin, and a growth factor cocktail containing 10 ng/ml of interleukins (IL)-3, (IL)-6, (IL)-7 and granulocyte colony-stimulating factor (G-CSF), 5 ng/ml of granulocyte macrophage colony-stimulating factor (GM-CSF), and 25 ng/ml of stem cell factor (SCF) (Peprotech).

### Cell lines

The leukemic DA1-3b cell line was generated by stable transfection of BCR-ABL1[14] [15] [16]. Isolation of tumor cells d60 and d365 have been described previously [14] [15] [16]. K562, KCL-22 and KU812 cell lines were grown in the same conditions. MS-5 mesenchymal cells were grown in α-MEM medium w/o nucleosides (Gibco) supplemented with 10% fetal calf serum, 50 U/ml penicillin, 50 mg/ml streptomycin, 2 mM L-glutamine and 2 mM pyruvate. The identity of HBL, LND, and Mel-4M was also confirmed by karyotyping and array comparative genomic hybridization testing.

### OCR and ECAR measurement, determination of cellular ATP

Extracellular acidification rate measurements (ECAR) and oxygen consumption rate (OCR) were measured using the Seahorse XFe24 analyzer (Seahorse Bioscience, Billerica, MA, USA). Detailed methods are provided in Supplementary Information files.

### Colony forming cell (CFC) assay

Clonogenic assay was realized with cells seeded into 35mm petri dish in semi-solid methylcellulose medium (Methocult^™^ M3231 for murin cells or Methocult^™^ H4230 for human cells, Stemcell Technologies). Cells were treated with the drugs for 72 hrs and centrifugated. Pellet was resuspended in Iscove’s modified Dulbecco medium (Lonza) supplemented with 2% fetal calf serum and 50 U/ml penicillin, 50 mg/ml streptomycin at 10 000 cells/ml, and cell suspension was added to methylcellulose medium (1000 cells/1ml/dish) and left at 37°C and 5% CO_2_. Colony forming efficiency was determined after 7 days using Leica DMI8 inverted microscope (Leica Microsystems) and quantified using Image J software.

### Metabolite flux

Two hundred thousand cells were supplemented with media containing uniformly labelled U-13C_6_ glucose (25 mM) or 13C-glutamine (2 mM) for 24 hrs. Detailed methods are provided in Supplementary Information files.

### Immunoblot analysis and Real-time quantitative reverse transcription

For immunoblot, cell lysates were prepared as described previously [17] Quantitative detection of mRNA was performed by real-time PCR using the Lightcycler 480 detector (Roche Applied Science, Manheim Germany) as previously published [17].

### Amino acid measurements

Amino acids concentration assay (μmol/l) was performed by high-performance liquid chromatography (Shimadzu C18 column, Kyoto, Japan) associated with tandem mass spectrometry (Sciex 3200 Qtrap, Framingham, MA) using the aTRAQ kit for amino acid analysis of physio-logical fluids (Sciex). Acquisition in the mass spectrometer was achieved by multiple reaction monitoring. Data recording and analysis were performed with Analyst software, v.1.6 (Sciex). Internal controls were systematically analyzed for each series of samples.

### In vivo Studies

The DA1-3b/C3H mouse model of tumor dormancy has been described previously [14] [15] [16]Seven-to eight-week-old C3H/HeOuJ female mice (Charles River Laboratories, Lyon, France) were intraperitoneally injected with 1 × 10^6^ DA1-3b, DA1-3b d60 or DA1-3b/d365. All animal experiments were approved by the Animal Care Ethical Committee CEEA.NPDC (agreement no. 2017022716306305).

### Statistical analysis

All data points are represented as means ± SD. Two-tailed Student’s t test was used to compare mean values between two groups. One-way or two-way analysis of variance (ANOVA) followed by Dunnett’s or Sidak post hoc testing was used to compare mean values between multiple groups. Statistical analysis was performed using Prism version 6.0f (GraphPad Software, La Jolla, CA). P < 0.05 were statistically significant.

## Results

### Metabolic organization of CML involves both glycolysis and glutamine dependent mitochondrial OXPHOS

We first compared the metabolism of DA1-3b leukemic cells expressing BCR-ABL1 to the isogenic cell line DA1 that does not express BCR-ABL1 (Fig. 1A-1C). There was a significant increase in both glycolysis (as judged by ECAR and gene expression) (Fig. 1A and 1B) as well as mitochondrial respiration (Fig. 1C) in cells transfected with p210 BCR-ABL1 compared to control cells. In BCR-ABL1-expressing DA1-3b cells, the mitochondrial respiration was largely sustained by glutamine (Fig. 1D) indicating that CML cells consume both glucose and glutamine. In the presence of glutamine, glucose and pyruvate, DA1-3b exhibited the highest proliferative rates (Fig. 1E). Under conditions of glucose or glutamine deprivation, DA1-3b cells were still able to proliferate although at a lower rate than in conditions with both nutrients (Fig. 1E). Glucose or glutamine starvation did not induce obvious increase in cell death but promoted G0/G1 cell cycle arrest (Fig. 1F). This result is consistent with the observation that inhibition of either glycolysis with 2DG or mitochondrial respiration with oligomycin A was insufficient to totally deplete DA1-3b (Fig. 1G, left panel) or primary CD34+ leukemic (Fig. 1G, right panel) cells in ATP. This is probably due to the development of compensatory mechanisms to prevent energy collapse. These results suggest that CML cells possess metabolic flexibility to survive for periods of carbon sources deprivation. Finally, only the inhibition of the two metabolic pathways depleted cells in ATP and was able to kill BCR-ABL1 expressing cells. All together, these results strongly suggest that only the depletion of carbon sources, glucose and glutamine, are required for energy crisis and subsequent CML killing.

**Figure 1.**
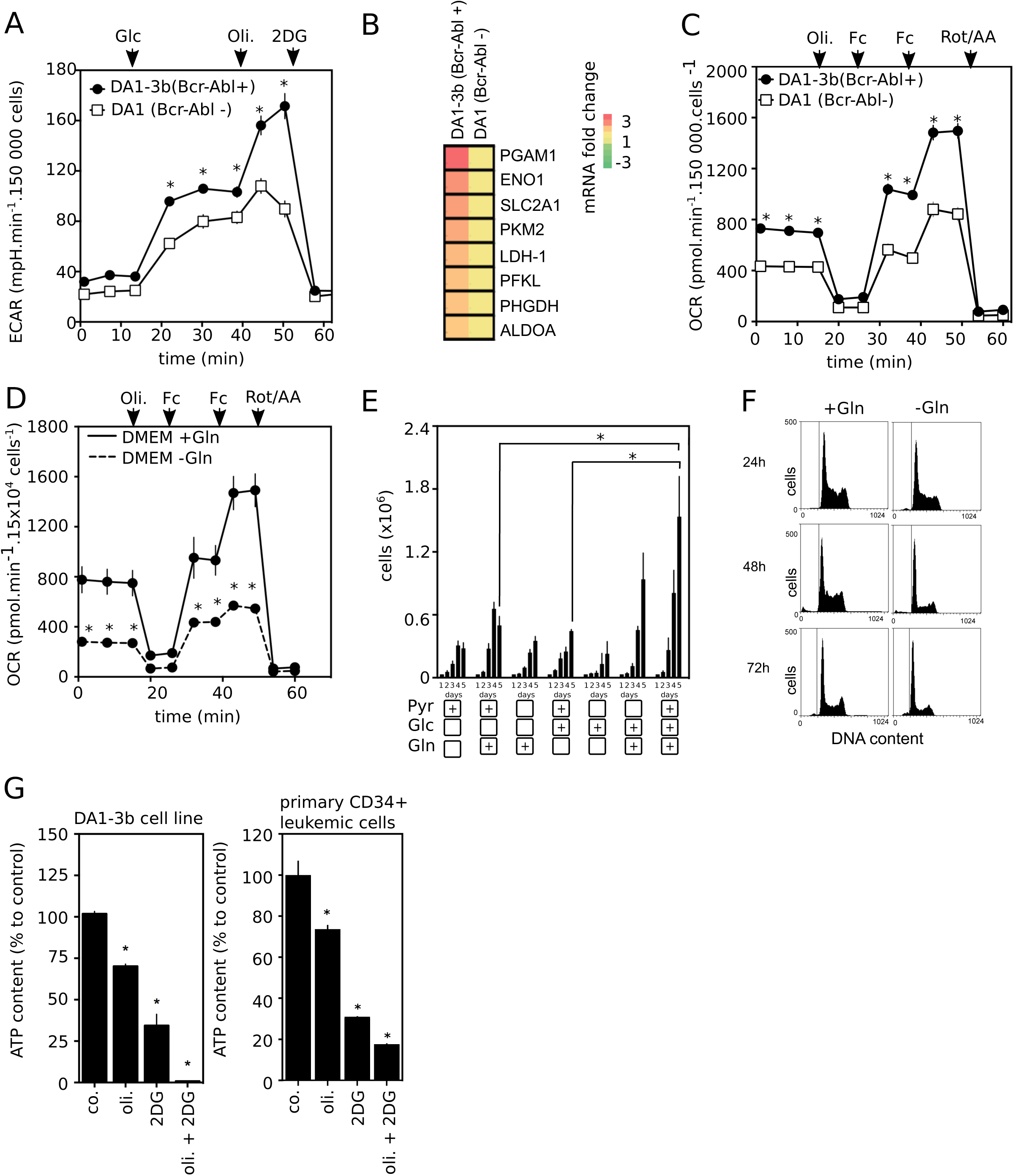
Glucose metabolism and mitochondrial respiration through glutamine oxidation are necessary for optimal metabolism and cell proliferation in CML cells. **(A)** Glycolytic activity and (C) oxygen consumption of BCR-ABL1+ (DA1-3b) and BCR-ABL1- (DA1) measured with Seahorse XFe24 extracellular flux analyzer after the injection of indicated drugs (Glc for glucose, Fc for FCCP, Oli for oligomycin A, 2DG for 2-deoxy-glucose, Rot/AA for rotenone and antimycin A) (n=3, * p < 0.05). (B) Glycolysis enzymes mRNA expression were quantified by RT-qPCR. 18S mRNA was used as housekeeping gene and data were expressed as mean of fold change (n=3, * p < 0.05). (D) DA1-3b leukemic cells were incubated in DMEM medium with glucose supplemented or not with glutamine for 24 hrs and oxygen consumption rate (OCR) was then measured. (E) DA1-3b cells were cultured in medium with or without glutamine, glucose or pyruvate as indicated. Proliferation was assessed by cell count from 1 to 5 days (mean +/-SD, n=3). (F) DA1-3b cells were cultured in medium with or without glutamine. After 24, 48 and 72 hrs, cell death was determined by measuring sub-G1 population using propidium iodide staining. Pictures are representative of three independent experiments. (G) DA1-3b cells (left panel) and primary CD34^+^ leukemic cells isolated from CML patient blood (right panel) were untreated (co.) or treated with oligomycin A (1µM), 2-DG (10 mM) and a combination of both molecules for 4 hrs. ATP content was measured by luminescence (mean +/- SD, n=3. * p < 0.05).

### Upon Imatinib exposure BCR-ABL1+ cells are dependent on mitochondrial metabolism for survival

Previous studies have shown that Imatinib displays a sustained inhibitory effect on glucose uptake and glycolysis through a reduction in the expression of key proteins such as GLUT-1 or PKM2 [9,18]. In the current study, we examined specifically the metabolism of leukemic cells that survive to Imatinib exposure at sub-lethal (Supplemental Fig S1A-D), and clinical relevant anti-proliferative concentrations [11]. At concentration below 0.2µM, Imatinib induced a strong antiproliferative effect in DA1-3b cells (Supplementary Fig S1A) but did not induce mitochondrial apoptosis as seen by the absence of translocation of phosphatidyl-serines, the maintain of high mitochondrial membrane potential (ΔΨm) and the low level of mitochondrial ROS (Supplementary Fig S1 B-D).

We performed metabolic flux analysis using 13C-labeled glucose in CML cells treated with vehicle or Imatinib for 24 hours before the incubation with labeled glucose. Cells treated with Imatinib displayed important impairment of glucose uptake and glycolysis as judged by the decrease in labeled glycolytic intermediates and labeled lactate (Fig. 2A). Moreover, it was accompanied by a reduction in the glucose flux through the nonoxidative pentose phosphate pathway (Fig. 2A). In agreement with the inhibition of glucose uptake and lactate production (supplementary Fig. S2A and S2B), we also observed a decrease in glycolysis-associated protein expression (supplementary Fig. S2C), as well as a decrease in ECAR in BCR-ABL1+ leukemic cells exposed to Imatinib (Fig. 2B) or to other clinical BCR-ABL1 inhibitors (Fig. 2C). Despite the decrease in glycolysis, Imatinib exposure did not deplete in the high-energy molecule, ATP indicating that cells that survive to Imatinib can maintain energy state in the absence of efficient glycolysis (Fig. 2D). This situation is compatible with the maintenance of mitochondrial metabolism in spite of Imatinib exposure, despite a slight decrease in basal OCR under TKI treatment (supplementary Fig. 2E). To explore whether glutamine was used to fuel mitochondrial activity in such conditions, we cultured cells in [13C5] L-glutamine in the presence or absence of Imatinib and analyzed intracellular metabolites by mass spectrometry. As shown in Figure 2E, cells that survive to Imatinib continued to use glutamine to produce the TCA intermediates alphaKetoGlutarate in the absence of efficient glycolysis (Fig. 2B). Thus, we observed that leukemic cells exposed to Imatinib relied on glutamine-dependent mitochondrial activity to survive. Indeed, these cells became highly sensitive to the withdrawal of glutamine but not asparagine (Fig. 2F). Thus, to survive to Imatinib, BCR-ABL1+ cells require glutamine-derived carbon that maintains the TCA cycle in the absence of glycolysis.

**Figure 2.**
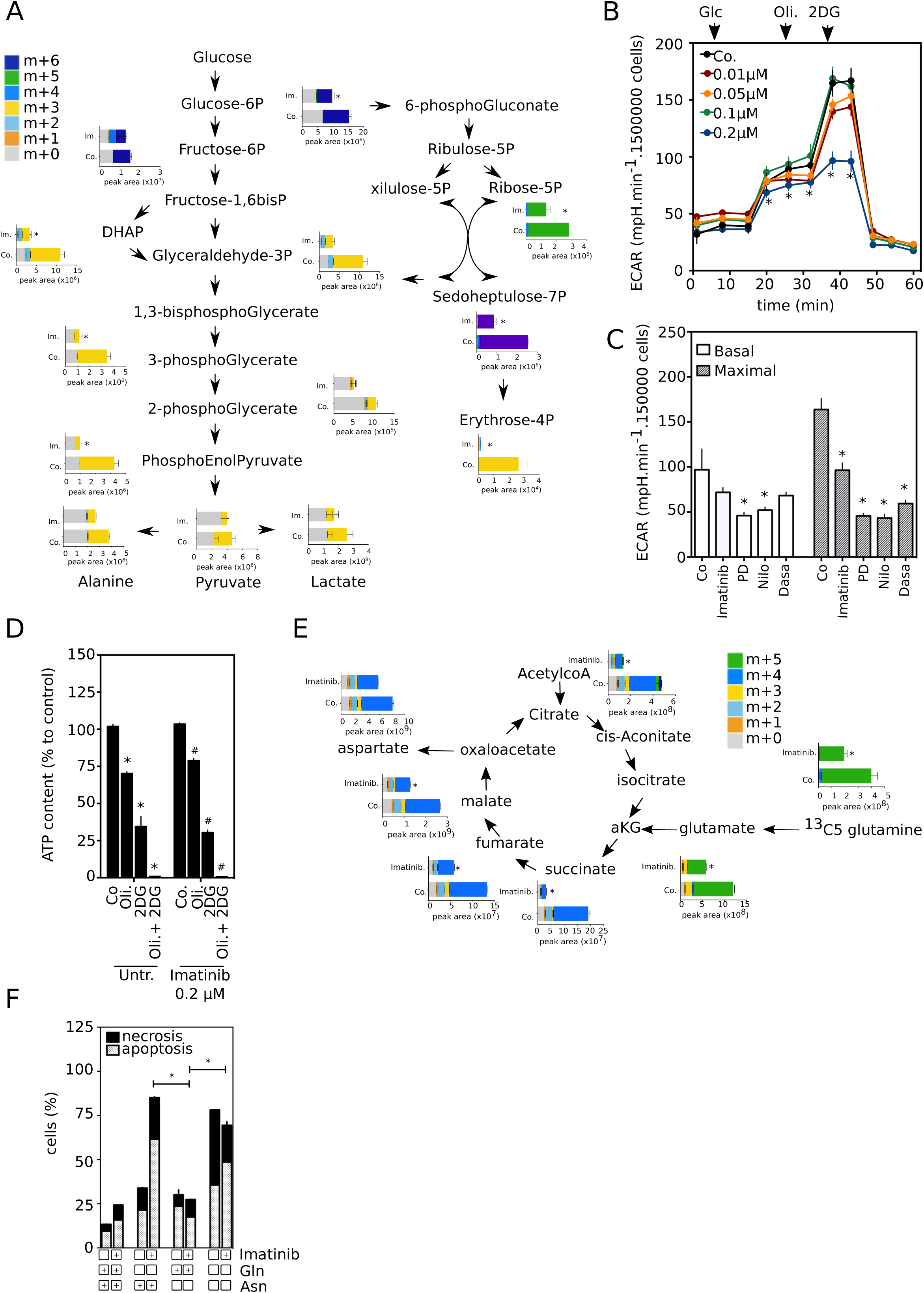
BCR-ABL1+ cells exhibit mitochondrial addiction and glutamine dependency under TKI exposure. (A) Isotopolog quantification of glycolysis intermediates have been measured by liquid chromatography-mass spectrometry analysis in DA1-3b cells treated with Imatinib 0.2µM (Im.) for 24 hrs or not treated (Co.) and grown in media containing U-13C_6_ glucose (mean ± SD, n=3, *p=0.05). The symbol ‘‘m+’’ indicates the number of carbon atoms of each metabolite labeled with 13C. (B) Glycolytic activity (ECAR) of DA1-3b cells treated with Imatinib (0.01-1 µM) for 24 hrs measured with Seahorse XFe24 extracellular flux analyzer after the injection of indicated drugs (Glc for glucose, Oli for oligomycin A, 2DG for 2-deoxy-glucose) (C) Variations of glycolytic activity (ECAR) was determined after addition of inhibitors (Glc for glucose, Oli for oligomycin A, 2DG for 2-deoxy-glucose) using Seahorse XFe24 extracellular flux analyzer (left panel). Leukemic cells were treated by sub-lethal concentration of TKI of BCR-ABL1 (Imatinib 0.2 µM, PD180970 0.01 µM, Nilotinib 5 nM, Dasatinib 2 nM) for 24 hrs, and basal and maximal ECAR were assessed. Basal glycolysis ans maximal glycolysis are measured following glucose or oligomycin injection respectively (mean +/-SD, n=3. * p < 0.05). (D) DA1-3b leukemic cells were treated with Imatinib (0.2 µM) for 72 hrs or with oligomycin A (1µM) or/and 2-DG (10mM) for 4 hrs. ATP levels were then measured by luminescence (mean +/-SD, n=3. * or # P < 0.05 respectively compared to control). (E) Isotopolog quantification of TCA cycle intermediates levels by liquid chromatography-mass spectrometry analysis in DA1-3b cells grown in media containing U-13C_5_ glutamine and treated or not with Imatinib (0.2 µM for 24 hrs) (mean ± SD, n=3, *p < 0.05). (F) DA1-3b cells were cultured in medium containing a combination as indicated of Imatinib (0.5 µM), glutamine (2 mM) and asparagine (4 mM) for 48 hrs. Necrosis and apoptosis were determined by flow cytometry analysis of Annexin V and Sytox blue staining (mean ± SD, n=3, *p < 0.05).

### L-asparaginases inhibit glutamine metabolism and reduce leukemic cell growth but are insufficient to eradicate BCR-ABL1+ cells

We have previously demonstrated that targeting glutamine metabolism has potential anti leukemic effects on myeloid leukemia cells [19,20]. Therefore, we sought to target glutamine addiction in Imatinib-surviving BCR-ABL1+ cells. L-asparaginases (*e*.*g*. the *E*.*coli-asparaginase*, Kidrolase) are used to treat pediatric and adult forms of acute lymphoblastic leukemia and are also used in pediatric AML. These therapeutically relevant components are able to deaminate L-asparagine into aspartate. These enzymes also deplete glutamine and the antileukemic activity correlates with their ability to deplete extra cellular asparagine better than glutamine at lower dose, and to deplete both amino acids at higher dose [21]. Consequently, we studied the metabolic effects of the FDA-approved Kidrolase in BCR-ABL1 + leukemic cells (Fig. 3). In conditions that deplete extracellular glutamine and asparagine (Fig. 3A-3B and Supplementary Fig. S3A), the flux experiment indicates that L-asparaginase depleted drastically the anaplerotic flux of glutamine into the TCA in BCR-ABL1+ cells (Fig. 3C). This was accompanied by a significant reduction in OCR (Fig. 3D) confirming that glutamine is a major energy source to fuel mitochondrial respiration in BCR-ABL1+ cells. As a result of mitochondrial inhibition, Kidrolase displayed strong antiproliferative effects without predominant cytotoxic activity in BCR-ABL1+ cells (Fig. 3E and supplementary Fig. S3B). This absence of important cell death in BCR-ABL1+ cells was compatible with the maintenance of high ATP level in Kidrolase-treated cells (Fig. 3F). Strikingly, L-asparaginase increased intracellular glycolytic intermediates in BCR-ABL1+ cells (Fig. 3G). Next, we tested whether the increase of glycolysis upon Kidrolase treatment allows leukemic cells to cope with metabolic stress. We found that the compensatory increase in glycolysis supported the survival of cells exposed to Kidrolase since inhibition of glycolysis with 2DG synergized with Kidrolase in inducing BCR-ABL1+ cell death (Fig. 3H). To determine whether the sensitivity to Kidrolase is a general feature of leukemic cells, we tested a large panel of BCR-ABL1 + or - leukemic myeloid cell lines. The antileukemic effects of Kidrolase were dose dependent and occurred in all tested cell lines. However, the antileukemic responses were highly heterogeneous in term of cell death observed up to 72h (supplementary Fig. S3C). Accordingly, we discovered an inverse correlation between glycolysis and Kidrolase-induced leukemic cell death (Fig. 3I). Thus, the FDA-approved glutamine inhibitor Kidrolase hinders glutamine-dependent mitochondrial metabolism but are insufficient to eradicate BCR-ABL1+ cells due to glycolytic compensation.

**Figure 3.**
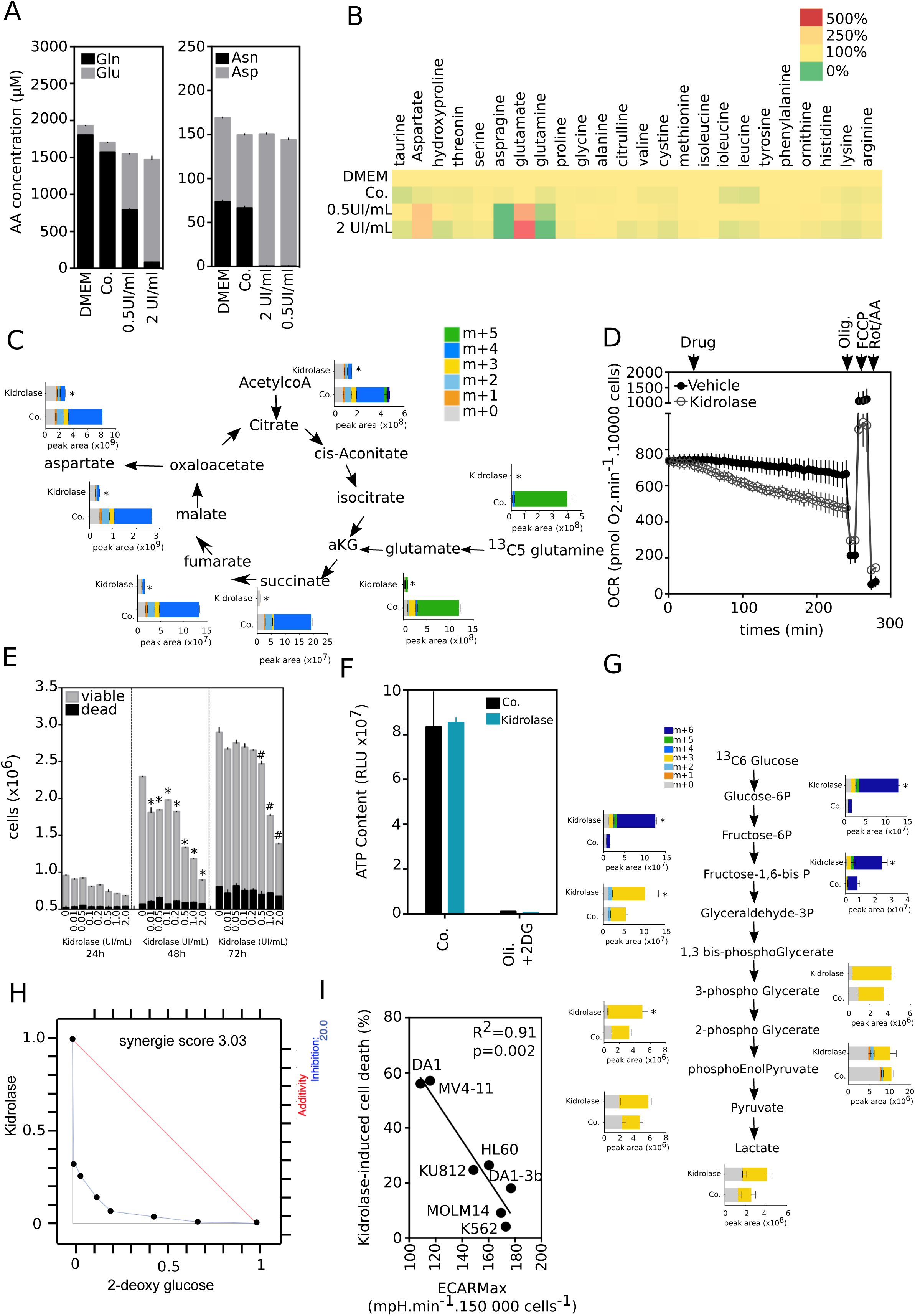
Myeloid leukemic cells are able to survive to Kidrolase-induced glutamine depletion through glycolysis. (A) and (B) DA1-3b cells were cultured in medium containing Kidrolase 0.5 or 2 UI/mL (Kid.) for 24 hrs. Cells were centrifugated, supernatent was removed and amino acids concentration as indicated was measured by HPLC/MS. (C) Isotopolog quantification of TCA cycle intermediates levels has been measured by LC-MS analysis in DA1-3b cells grown in media containing U-13C_5_ glutamine and treated or not with Kidrolase (0.5 UI/mL) for 24 hrs (mean ± SD, n=3, *p=0.05). The symbol ‘‘m+’’ indicates the number of carbon atoms of each metabolite labeled with 13C. (D) Assessment of mitochondrial respiration in DA1-3b cells by measuring OCR with XF24e Seahorse. The following molecules have been injected subsequently: drug (Kidrolase or vehicle control), oligomycin (Olig.), FCCP and rotenone/antimycine A (Rot/AA). (E) Cell proliferation and cell death (by PI staining) in DA1-3b cells treated by Kidrolase (from 0 to 2 UI/mL) were determined at 24, 48 and 72 hrs (mean ± SD, n = 3, * or # p < 0.05 respectively compared to control). (F) Intracellular ATP level of DA1-3b cells exposed to oligomycin (oli.), 2-deoxy-glucose (2DG) or a combination of the two inhibitors for 4 hrs, with or without 72 hrs pre-incubation with Kidrolase 1 UI/mL (mean ± SD, n=3, *p=0.05). (G) Isotopolog quantification of glycolysis intermediates measured by LC-MS analysis in DA1-3b cells treated with Kidrolase 0.5 UI/mL for 24 hrs and grown in media containing U-13C_6_ glucose (mean ± SD, n=3, *p=0.05). (H) DA1-3b cells were cultured with combination of 2-DG (0.033 – 100 mM) and Kidrolase (0.00033 – 10 UI/mL) for 72 hrs and cell proliferation inhibition was quantified by fluorescence using CyQUANT Cell Proliferation Assay. The response of the combination was compared with its single agents against the widely used Loewe model for drug-with-itself dose additivity using Chalice software and presented as an isobologram. (mean ± SD, n=3, *p=0.05). (I) Correlation between maximal glycolytic activity deterrmined by XFe24 Seahorse after glucose/oligomycin injection (ECAR Max) and cell death (determined by PI staining) induced by 48 hrs Kidrolase treatment (0.5 UI/mL) in 8 myeloid leukemic cell lines (p =0.002, R^2^=0.91).

### Dual treatment with Imatinib and Kidrolase significantly induces death of BCR-ABL1+ leukemic cells

Given the metabolic flexibility of BCR-ABL1+ cells observed above, we hypothesized that the combination of Imatinib and L-asparaginases such as Kidrolase could be of therapeutical interest blocking both glycolysis and mitochondrial metabolism. Metabolic flux analyses indicate that cells treated with the combination of Imatinib and Kidrolase caused much more impairment of the carbon flux into the TCA cycle, than either drug alone (Fig.4A right panel). This decrease of TCA activity was not linked to modification of respiratory chain protein expression as seen by immunoblot of several proteins of each complexe (supplementary Fig. 3E). The combination of drugs also impeded the glycolytic flux with efficiency (Fig. 4A left panel). In BCR-ABL1+ cell lines, Kidrolase synergized with Imatinib to induce cell death (Fig. 4B-4C and supplementary 3F). The combination of Imatinib with Kidrolase enhanced cell death by potentiating the intrinsic pathway of apoptosis through the downregulation of Bcl-2 and Bcl-XL protein levels and Bim upregulation (Fig. 4D). The combination of Kidrolase and Imatinib exhibited the most pronounced killing effect in comparison to the association of Kidrolase and other anti-leukemic drugs (daunorubicin or idarubicin) (Fig. 4E). Kidrolase acted in synergy with Imatinib in BCR-ABL1 + leukemia cells even when cells were cultured on the MS5 stroma cell line (Fig. 4F) or in hypoxic conditions (Fig. 4G), situations known to protect leukemia cells from the effects of L-asparaginase [22] or to Imatinib [23]. These results suggest that Imatinib plus Kidrolase elicit death synergistically in BCR-ABL1 + leukemic cells.

**Figure 4.**
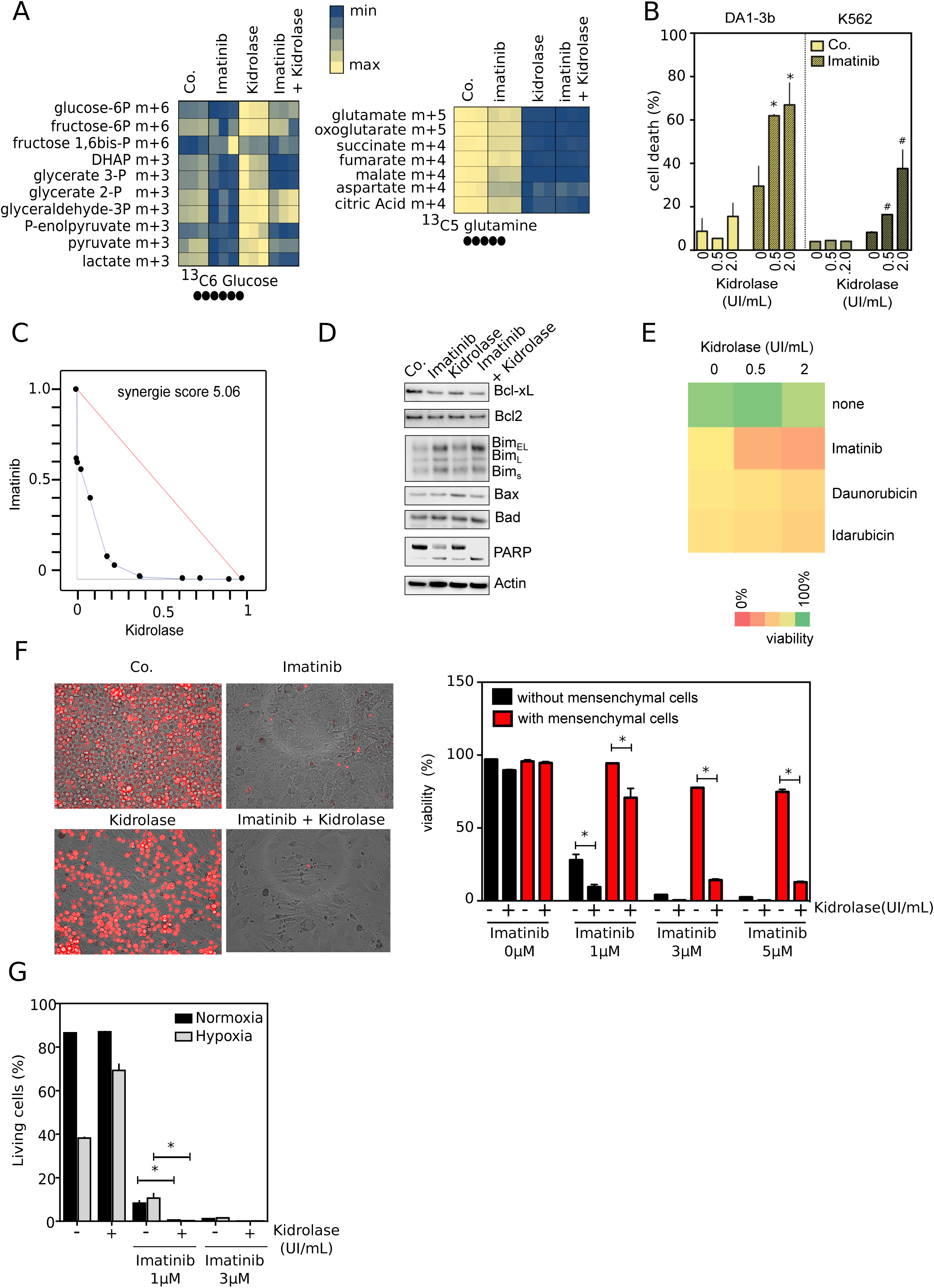
Imatinib and Kidrolase drug combination is effective to target glycolysis and mitochondrial metabolism and to induce cytotoxic effects. (A) Isotopolog quantification of glycolysis (left panel) and TCA cycle (right panel) intermediates calculated as percentage of the total metabolite pool following liquid chromatography-mass spectrometry in DA1-3b cells treated with Imatinib (0.2 µM) and Kidrolase (0.5 UI/mL) for 24 hrs and grown in media containing U-13C_6_ glucose (left panel) or U-13C_5_ glutamine (right panel) (n=3). The symbol ‘‘m+’’ indicates the number of carbon atoms of each metabolite labeled with 13C. (B) Determination of DA1-3b and K562 cell death after 48hrs exposure to Imatinib (0.5 µM) and Kidrolase treatments as indicated (mean ± SD, n = 3, * or #p < 0.05 respectively compared to control). (C) DA1-3b cells were cultured with combination of Imatinib (0.0001 – 100 µM) and Kidrolase (0.00033 – 10 UI/mL) for 72 hrs and cell proliferation inhibition was quantified by fluorescence using CyQUANT Cell Proliferation Assay). Isobologram have been determined as seen in Fig. 3. (D) Immunoblotting of pro- and anti-apoptotic proteins (as indicated) in DA1-3b cells treated with Imatinib (0.5 µM), Kidrolase (2 UI/mL) or a combination of both drugs for 24 hrs. Pictures are representative of three independent experiments. Actin was used as loading control. (E) DA1-3b cells were treated with a combination of Kidrolase and anticancer drugs (Imatinib 0.5 µM, Daunorubicin 0.01 µM, Idarubicin 0.001µM) for 48 hrs and viability was assessed by cytometric analysis of annexin V and Sytox stainings. (F) Phase-contrast analysis of DA1-3b cells transfected with DsRed co-cultured with MS-5 mesenchymal cells and treated by Imatinib (3 µM) and Kidrolase (2 UI/mL) for 7 days (left panel). Viablity of DA1-3b in mono-culture or in co-culture with mesenchymal cells after 48hrs treatments with Imatinib (1-5 µM) and Kidrolase (2 UI/mL) (right panel). Qauntification was expressed as mean ± SD (n=3, *p=0.05). (G) DA1-3b cell lines were exposed to Imatinib and Kidrolase treatments in normoxia (20% O_2_) or hypoxia (1% O_2_) for 72 hrs. Viability was assessed by flow cytometry after PI stainings (mean ± SD, n = 3, * p < 0.05).

### Imatinib synergized with Kidrolase to eradicate BCR-ABL1+ persistent stem cells and leukemia-initiating cells in CML patients

Next, we explored the potential of the synergistic combination of Imatinib plus Kidrolase against subpopulations of short-term relapse-inducing CML cells. First, the subpopulation of cells that survived in the presence of Imatinib for 7 days was enriched in progenitors immature CD34+ CD38-progenitors (Fig. 5A) and withdrawal of Imatinib lead to proliferation of blast cell clones (Fig 5B).

**Figure 5.**
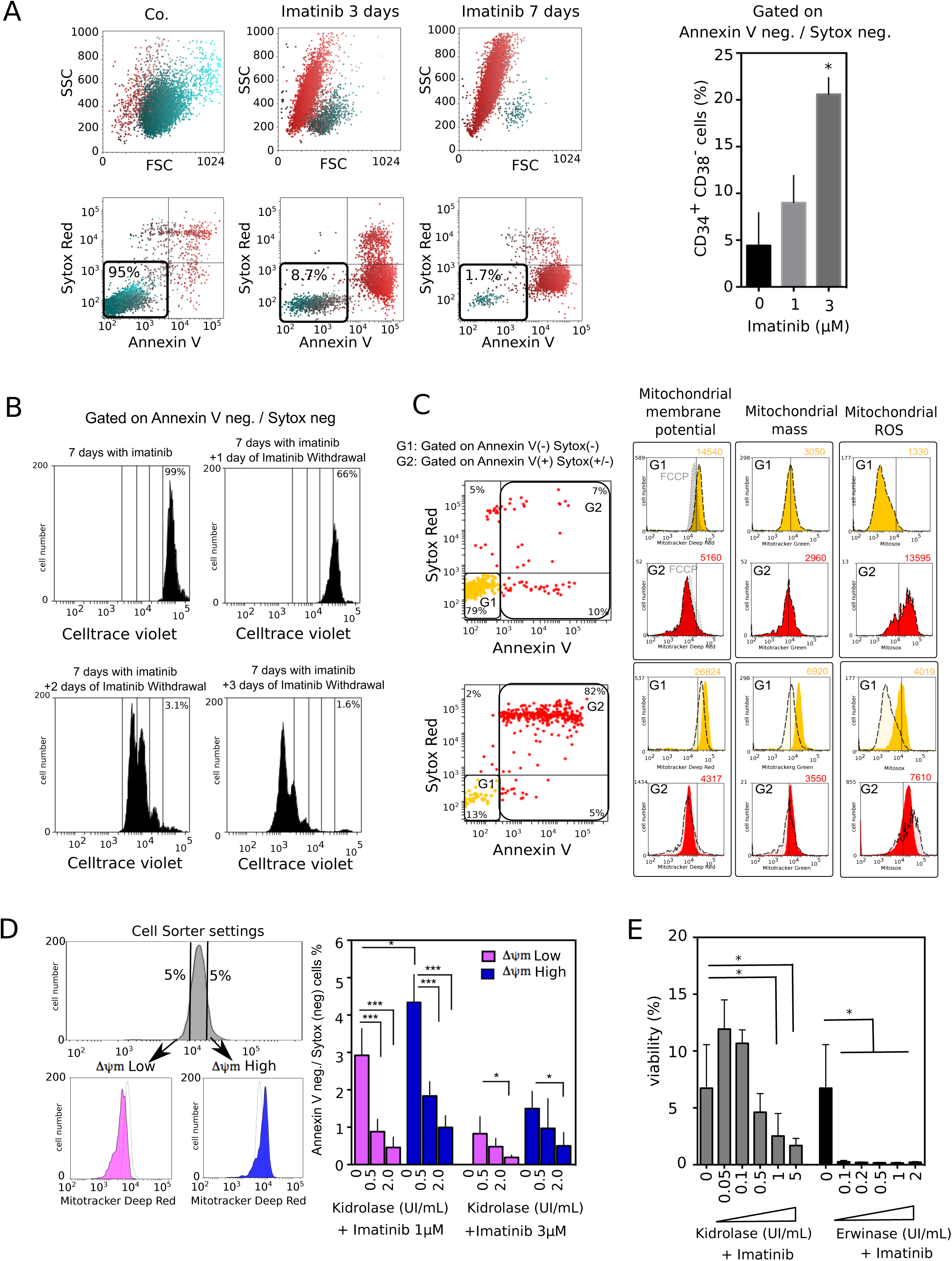
TKI and Kidrolase combination reduces Bcr Abl persistant cells following TKI treatment. (A) DA1-3b cells have been treated by Imatinib and viability has been assessed by cytometric analysis using Annexin V and sytox blue stainings. The percentages of CD34+ CD38-cells have been determined in persistant cells subpopulation (Annexin V neg / Sytox neg) following 7 days of treatment. (mean ± SD, n = 3, * p < 0.05). (B) DA1-3b cells have been treated by Imatinib for 7 days and medium has beeen changed for Imatinib-free medium (Imatinib withdrawal condition) containing CellTrace Violet probe. CellTrace Violet fluorescences have been measured by flow cytometry following 1, 2 and 3 days after Imatinib withdrawal. (C) DA1-3b cells were untreated or treated by Imatinib (0.5 µM) for 7 days. Mitochondrial membrane potential, mitochondrial mass and mitochondrial ROS production were measured in living (yellow histograms) and dead (red histograms) cells using flow cytometry analysis of mitotracker deep red, mitotracker green and mitosox red stainings respectively, and annexin V and sytox stainings. For membrane potential analysis, FCCP was used as positive control for depolarization of mitochondrial membrane (grey histogram). The mean of fluorescence is indicated at the top right corner of each histogram. Cytofluorimetric profiles are representative of three independent experiments. (D) Using cell sorting by flow cytometry, DA1-3b cells have been separated in two subpopulations characterized by low or high mitochondrial membrane potential (ΔΨm) in comparison with unsorted population. Both populations have been treated with Imatinib (at the indicated concentrations) for 7 days and the percentage of persistant cells (annexin V neg. / sytox neg) have been determined by flow cytometry. (mean ± SD, n = 3, * p < 0.05; *** p < 0.005). (E) Cell death (by PI staining) in DA1-3b cells treated by Kidrolase or Erwinase (from 0 to 2 UI/mL) in combination with Imatinib (0.5uM) was determined after 7 days (mean ± SD, n = 3, *p < 0.05).

This persistent population which survived during TKI expsoure presented high level of mitochondrial metabolism as determined by their mitochondrial potential, mitochondrial mass and the overproduction of mitochondrial ROS (Fig. 5C). To confirm the predominant role of mitochondrial metabolism, we isolated two populations characterized by low (ΔΨ_m_ low) or high mitochondrial (ΔΨ_m_ high) membrane potential using cell sorting. As expected, the percentage of persistent leukemia cells was higher in ΔΨ_m_ high DA1-3b cells in comparison with ΔΨ_m_low cells, whereas the combination of Imatinib and Kidrolase is efficient to eradicate both subpopulations. Finally, we showed that Erwinase, another L-asparaginase, also synergized also with Imatinib to kill persistent BCR-ABL1+ cells (Fig 5E).

Next, we studied the effect of this drug association on long-term persistent BCR-ABL1+ cells originated from a mouse model of *in vivo* leukemia dormancy [15]. In our model, the DA1-3b/d60 and DA1-3b/d365 cells were derived from the BCR-ABL1 + DA1-3b cells injected in mice and isolated after 2 months or 1 year of tumor dormancy, respectively [24] (Fig. 6A). DA1-3b/d60 and DA1-3b/d365 were injected intraperitoneally in mice to developp a lethal leukemia (Fig 6B). As expected, death of DA1-3b/60 and DA1-3b/d365-bearing mice was delayed in comparison with DA1-3b WT mice, confirming the maintenance *in vivo* of the dormant phenotype (Fig 6B). Persistent-leukemic DA1-3b/d60 and DA1-3b/d365 cells were completely refractory to Kidrolase monotherapy and also partially resistant to treatment with Imatinib (Fig. 6C and 6D). However, the combination of both two drugs eradicated long-term persistent cells as judged by the induction of apoptosis (annexin V staining) (Fig 6D) and the loss of colony-forming potential (Fig 6E). Indeed, Kidrolase in combination with Imatinib reduced colony formation of DA1-3b cells by more than 90 % compared to Imatinib alone (Fig. 5E). Then, we assessed the *ex vivo* effects of Kidrolase plus Imatinib in different CD34+ and CD38-subpopulations of primary CML cells from patients at diagnosis (n=2; Fig. 6F) and in primary CML CD34+ progenitor cells from newly diagnosed patients (n=7; Fig. 6G). We show that the killing effect was more pronounced on the stem cell-enriched CD34+ CD38-subpopulation as compared to more differentiated CML cells (Fig. 6F). Furthermore, the combination of both drugs was effective to significantly decrease viability of CD34+ cells from newly diagnosed patients compared to individual treatment (Fig. 6G). Importantly, and as expected, the combination had no effect on CD34+ progenitor cells from healthy individuals (n=3; Fig. 6F).

**Figure 6.**
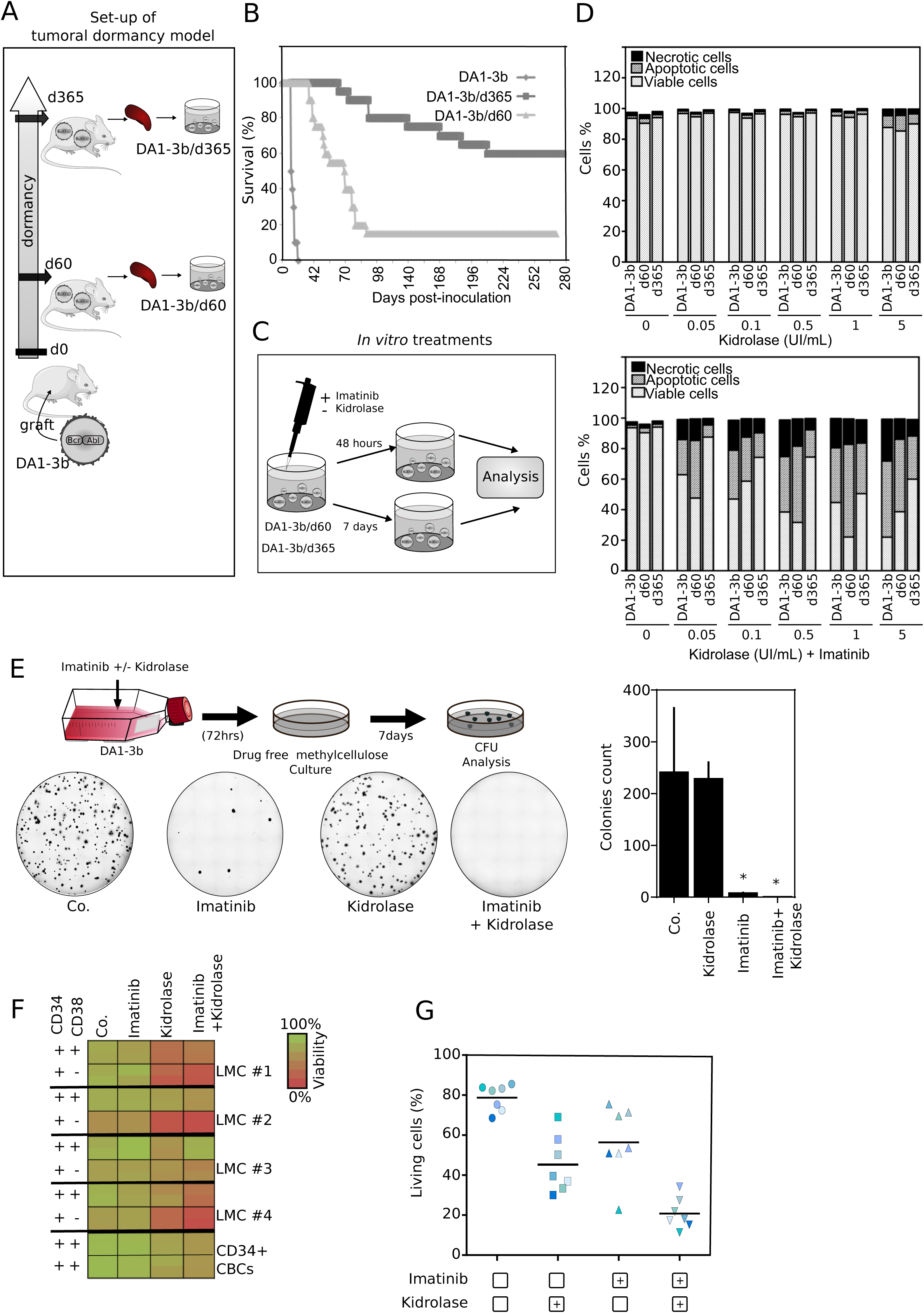
TKI and Kidrolase combination increases chemosensitivity in different models of TKI resistance. (A) Schematic representation of tumor dormancy model set up and treatments on leukemic dormant cells d60 and d365. Previously, mice were immunized with irradiated IL12–transduced DA1-3b cells, challenged with wild-type DA1-3b cells and randomly killed after 60 days or 365 days follow-up [24]. Leukemic residual cells were collected from bone marrow of sacrificed mice and used to generate stable cell lines (DA1-3b/d60 or DA1-3b/d365). (B) Lethal leukemia developped in mice injected intraperitoneally with DA1-3b wt, DA1-3b/d60 or DA1-3b/d365 cells. (C-D) DA1-3b, DA1-3b/d60 and DA1-3b/d365 cells were treated by Imatinib and Kidrolase for 48 hrs and cell death was assessed using flow cytometry analysis of annexin V and PI stainings. (n=3). (E) Schematic representation of colony forming cell (CFC) assay set up (see materials and methods for more details) (upper panel). DA1-3b cells were treated *in vitro* with Imatinib 1 µM (Ima.) and/or Kidrolase 2 UI/mL (Kidro) for 72 hrs. Cells were then seeded in semi-solid medium and left to incubate for 7 days. Images representative of 3 experiments are shown (lower panel). The clonogenic potential of leukemic cells after undergoing treatments are measured by colonies count (right panel) (mean ± SD, n = 3, *p < 0.05). (F) Assessment of CD34+ CD38+/- patient primary leukemic cell sub-population viability (PI assay) from CML patients (n=2) and CD34^+^ progenitors from cord blood cells (CBCs) (n=3), after 48 hrs exposure to Imatinib (3 µM) and Kidrolase (2 UI/mL). (G) Viability of CD34^+^ cells isolated from CML patient cytapheresis treated with Imatinib and Kidrolase for 48 hrs (n=7).

Thus, the combined use of Imatinib and Kidrolase synergistically increase selective cytotoxicity in CML stem cells.

## Discussion

We have characterized the metabolic effects of the FDA-approved drug Kidrolase in association with TKI in myeloid leukemia and demonstrated that they displayed synergistic antimetabolic effects that eradicate CML stem cells. It is well established that TKI such as Imatinib possess potent anti-Warburg effect. Imatinib efficiently hampers glucose metabolism through the reduction of GLUT-1 surface localization [9], inhibition of BCR-ABL1-mediated PKM2 phosphorylation or the modulation of PKM isoforms [18]. However, inhibition of glycolysis alone is often insufficient to eradicate cells due to compensatory activating metabolic pathways [6]. Upon Imatinib exposure, mitochondrial oxidative metabolism is maintained at high levels due to fatty acid [18] and/or glutamine oxidation (Fig 2). This suggests that remaining TKI-tolerant cells become addicted to mitochondrial activity for survival [13]. In agreement with our results, mitochondrial metabolism is spared by FLT3^ITD^ TK inhibitors in myeloid cells [25]. The mechanisms that result from a glycolytic metabolism shift toward oxidative metabolism remain largely unknown. However, we and others [25] have observed a significant increase in mitochondrial mass upon TKI exposure suggesting important changes in mitochondrial biogenesis. As a consequence of mitochondrial addiction, TKI-tolerant cells were highly sensitive to the antileukemia effects of mitochondrial targeting drugs. Myeloid leukemia present numerous mitochondrial-druggable targets [26]. Thus, the CPT1 (carnitine O-palmitoyltransferase 1) inhibitors that reduce fatty acid oxidation and mitochondrial OXPHOS decreased significantly the number of quiescent leukemia progenitor cells [27]. The combination of mitochondrial drugs and TKI consists in a rational approach that considers complementary mechanisms of action as the therapeutic aim. TKI and mitochondrial inhibitors address the two compensatory aspects of the metabolism, glycolysis and mitochondrial oxidation that none of the monotherapy can achieve alone to kill leukemia cells. Several preclinical studies evidenced that TKI, such as inhibitors of FLT3^ITD^ can synergize with mitochondrial inhibitors leading to potent antileukemia effects [25,28-30]. Interestingly, inhibition of the mitochondrial “booster” STAT3, [31] or of mitochondrial translation [13] synergize with BCR-ABL1 inhibitors in CML. Here, we completed the frame including the inhibition of glutamine-dependent mitochondrial metabolism. Genetic and pharmacological inhibition of the first enzyme in glutamine metabolism is synthetically lethal with FLT3-ITD-TKI in myeloid leukemia cells [25]. This effect seems related to the depletion of glutathione, enhancement of mitochondrial oxidative stress resulting in leukemia cell death [29]. We have used L-asparaginase to deplete extracellular glutamine and therefore mitochondrial metabolism. To best translate *in vitro* results into clinical application, we have chosen the existing FDA-approved drug Kidrolase. However, L-asparaginase use in the clinic is associated with a number of dose-dependent toxicities [32]. When combined with TKI, L-asparaginase should be less toxic since exposure to BCR-ABL1 inhibitors specifically increases the sensitivity of BCR-ABL1+ cells to mitochondrial targeting drugs (results and [30]). As demonstrated (Fig. 3A), L-asparaginase depleted the extracellular AA, glutamine and asparagine. Several studies uncovered a crucial role of asparagine in cancer cell growth and proliferation [33] as well as cell survival to glutamine deprivation [34]. Thus, asparagine is required for anabolism and cell proliferation in the absence of glutamine [35]. For these reasons, the synergistic effect of L-Asparaginase observed here is probably based on the depletion of both extracellular asparagine and glutamine. It is noteworthy that LSCs relies on AA including glutamine and glutamate to maintain OXPHOS for survival [36].

Targeting mitochondrial metabolism has been proposed as a therapeutic approach against CML stem cells [13,37,38]. Recently a therapeutic drug combination targeting mitochondrial metabolism has demonstrated efficacy against LSCs in patients with AML [39]. Accordingly, we have shown that both glycolysis and glutamine-dependent mitochondrial metabolism had to be impaired to eradicate LSCs. Targeting compensatory pathways of glutamine metabolism in CML stem cells can improve the efficacy of cancer treatments that impair glucose utilization. Thus, we have provided pre-clinical evidence that the antimetabolic cooperativity by the combination of oncogene tyrosine kinase inhibitors and mitochondrial inhibitors constitutes a novel interesting therapeutic approach to eradicate LSC. This antimetabolic strategy is able to target CML stem cells and could therefore limit therapeutic failure in patients following the discontinuation of TKI therapy.

## Acknowledgements

This work received a financial support from INSERM, UNIVERSITE DE LILLE, Ligue contre le Cancer (to PM and JK), a special financial support from the Association pour l’Etude des Anomalies Congénitales Neurodev of Pr. B. Poupard (to PM). AT is a recipient of a CHRU Lille-Région Nord-Pas de Calais fellowship. RK and QF are recipients of a University of Lille fellowship.

## Abbreviations

CML: (chronic myeloid leukemia)
TKI: tyrosine kinase inhibitors)
BCR-ABL1: B Cell Receptor-Abelson)
LSC: leukemic stem cell)

