## Supplemental information for "Antimetabolic cooperativity with the clinically approved kidrolase and tyrosine kinase inhibitors to eradicate cml stem cells"

Supplementary information

Materials and Methods

*OCR and ECAR measurement, determination of cellular ATP*

Extracellular acidification rate measurements (ECAR) and oxygen consumption rate (OCR) were measured using the Seahorse XFe24 analyzer (Seahorse Bioscience, Billerica, MA, USA). Detailed methods are provided in Supplementary Information files. Seahorse XF24 microplates were pre-coated with Corning^TM^ Cell-Tak (Fisher Scientific). Cells were suspended in medium containing DMEM (D5030, Sigma-Aldrich) with L-glutamine (2mM) and NaCl (32 mM) for glycolysis, or L-glutamine (2mM), glucose (10 mM) and pyruvate (1 mM) for oxygen consumption, and seeded at 150 000 cells/100µl/well for cell lines and 250 000 cells/100µl/well for primary cells. Cells were left to adhere by two successive centrifugations at low speed (650 rpm, then 450 rpm), and the microplate was left to stabilized for at least 20 min at 37°C in CO_2_-free incubator, after that 400 µl of warm medium were added to each well. Measurement of ECAR was done at baseline and after injections of the following molecules: D-glucose (10 mM), oligomycin A (1 µM), an ATP synthase inhibitor, and 2-deoxy-glucose (2-DG, 10 mM), a competitive inhibitor of glucose. For OCR measurement, the following molecules were added: oligomycin A (1µM), FCCP (0.25-0.5 µM), rotenone (1 µM) and antimycin A (1 µM).

For cellular ATP measurement, cells were seeded at 20 000 cells/100µl for cell lines or 100 000 cells/100µl for primary cells. After indicated time of treatments, Celltiter Glow Assay kit (Promega) was used according to the manufacturer’s procedures.

Cell proliferation was determined using cell counter Beckman cell counter or by cell numbers recorded after being seeded, using CyQuant direct proliferation kit from Invitrogen (Carlsbad, CA, USA).

*Cytofluorometric analysis*

Evaluation of cell viability was performed by determining the percentage of sub-G1 cells with propidium iodide staining (50 μg per ml, 30 min, 4°C) (P4864, Sigma-Aldrich, St. Louis, MO, USA) or by annexin V-FITC (0.45 µg per sample, 10 min, RT) (Biolegend), annexin V-APC (0.45 µg per sample, 10 min) (Biolegend), sytox blue (1µM, 10 min, RT) (Thermo Fisher Scientific), sytox red (5 nM, 10 min, RT) (Thermo Fisher Scientific), Cell trace violet (1µM, 15min, RT) (Thermo Fischer Scientific).

The CD34/CD38 cell profile was analyzed in one single tube containing the following molecules: for human cells CD34 ECD (clone 581, Beckman Coulter), CD38 APC (clone HIT2, Biolegend), CD45 BUV395 (clone HI30, BD Biosciences), for murin cells CD34 PE (clone MEC14.7, Biolegend), CD38 APC (clone 90, Biolegend), CD45.2 Brilliant violet 421 (clone 104, Biolegend) or a control isotype antibody.

Mitochondrial content, mitochondrial membrane potential and mitochondrial ROS production were assessed by MitoTracker Green (125 nM, 30 min, 37°C) (Thermo Fisher Scientific), MitoTracker Deep Red (125 nM, 30 min, 37°C) (Thermo Fisher Scientific) and Mitosox (2.5 µM, 30 min, 37°C) (Thermo Fisher Scientific) stainings respectively.

Fluorescence levels following cellular staining were analyzed by FACS LSR Fortessa X20 (Becton Dickinson).

*Immunoblot analysis*

Cell lysates were prepared as described previously [18] then 20 μg proteins were separated on a 4–12% SDS- PAGE and transferred to nitrocellulose membrane. After blocking for 1 h in 10% milk or Bovin Serum Albumine in TBS Tween buffer or according to manufacturer’s recommandations, membranes were then probed with antibodies (1:1000 for primary antibodies and 1:2000 for secondary antibodies) as described in Supplementary Table 1. Horseradish peroxidase- conjugated secondary antibodies from Rockland Immunochemicals Inc. (Limerick, PA) were used at 1:2000 for 1 h then detection was carried out by enhanced chemoluminescence.

*Real-time quantitative reverse transcription*

Quantitative detection of mRNA was performed by real-time PCR using the Lightcycler 480 detector (Roche Applied Science, Manheim Germany) as previously published [18]. The transcript levels in triplicates were compared using the Pfall method after normalization to those of α-tubulin.

*Glucose and lactate measurements*

Glucose and lactate were measured in the extracellular medium using a SYNCHRON LX20 Clinical system (Beckman Coulter, Fullerton, CA USA).

*Metabolite flux*

Two hundred thousand cells were supplemented with media containing uniformly labelled U-13C_6_ glucose (25 mM) or 13C-glutamine (2 mM) for 24 hrs. Cells were rinsed with ice-cold 0.9% NaCl solution and metabolites were extracted by adding 250 µl of a 50/30/20 solution (methanol-acetonitrile 10 mM Tris-HCl pH 9.4) to the cells. Plates were then incubated for 3 min on ice. Cells were transferred to an eppendorf tube before centrifugation at 20 000g for 10 min at 4°C. Supernatants were then transferred to a fresh tube and stored at -80°C, and pellets were used for BCA/protein assay to normalize each condition to the same cell number. Separation of metabolites prior to Mass Spectrometry (MS) measurement was performed using a Dionex UltiMate 3000 LC System (Thermo Scientific) coupled to a Q Exactive Orbitrap mass spectrometer (Thermo Scientific) operating in negative ion mode. Practically, 15 μl of the cellular extract was injected on a C18 column (Aquility UPLC®HSS T3 1.8µm 2.1x100mm) and the following gradient was performed by solvent A (H2O, 10mM Tributyl-Amine, 15mM acetic acid) and solvent B (100% Methanol). Chromatographic separation was achieved with a flowrate of 0.250ml/min and the following gradient elution profile: 0min, 0%B; 2min, 0%B; 7min, 37%B; 14min, 41%B; 26min, 100%B; 30min, 100%B; 31min, 0%B; 40min, 0%B. The column was placed at 40°C throughout the analysis. The MS operated both in full scan mode (m/z range: 70-1050) using a spray voltage of 4.9 kV, capillary temperature of 320°C, sheath gas at 50.0, auxiliary gas at 10.0. The AGC target was set at 3e6 using a resolution of 140 000, with a maximum IT fill time of 512 ms. Data collection was performed using the Xcalibur software (Thermo Scientific).
