## Supplementary figures and images for "Antimetabolic cooperativity with the clinically approved kidrolase and tyrosine kinase inhibitors to eradicate cml stem cells"

### Supplemental figure 1

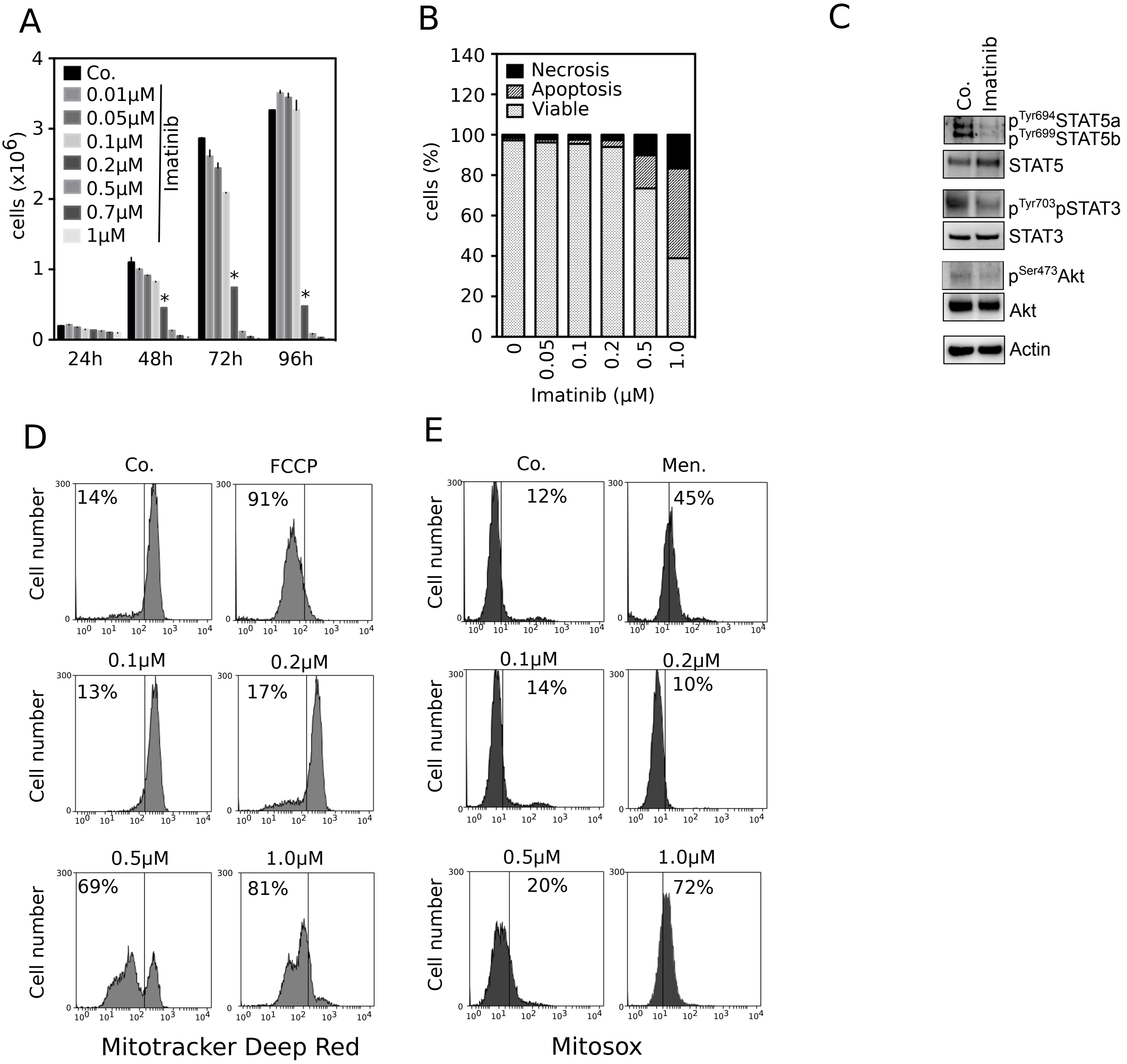


Supplementary Figure 1

### Supplemental figure 2

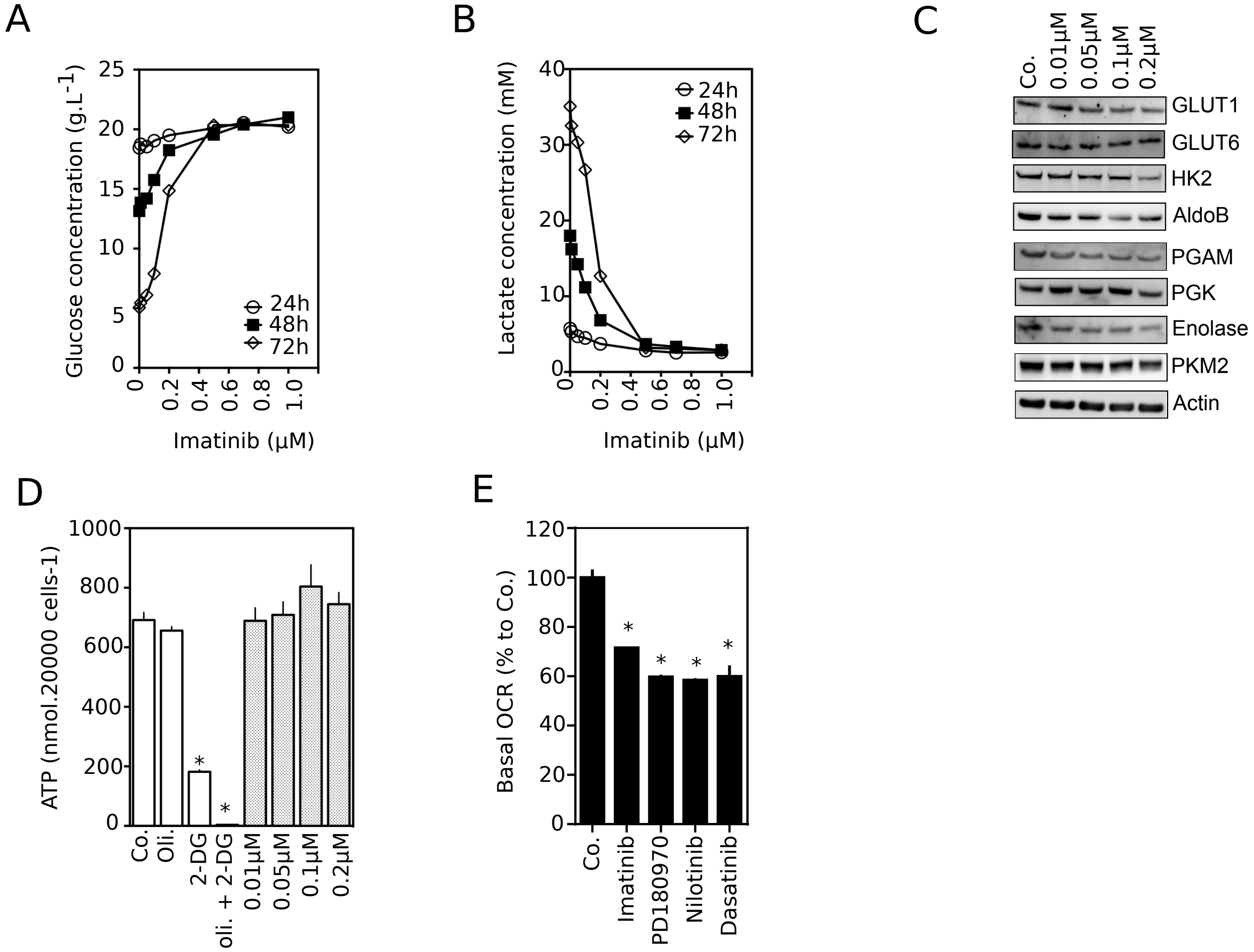


Supplementary Figure 2

### Supplemental figure 3

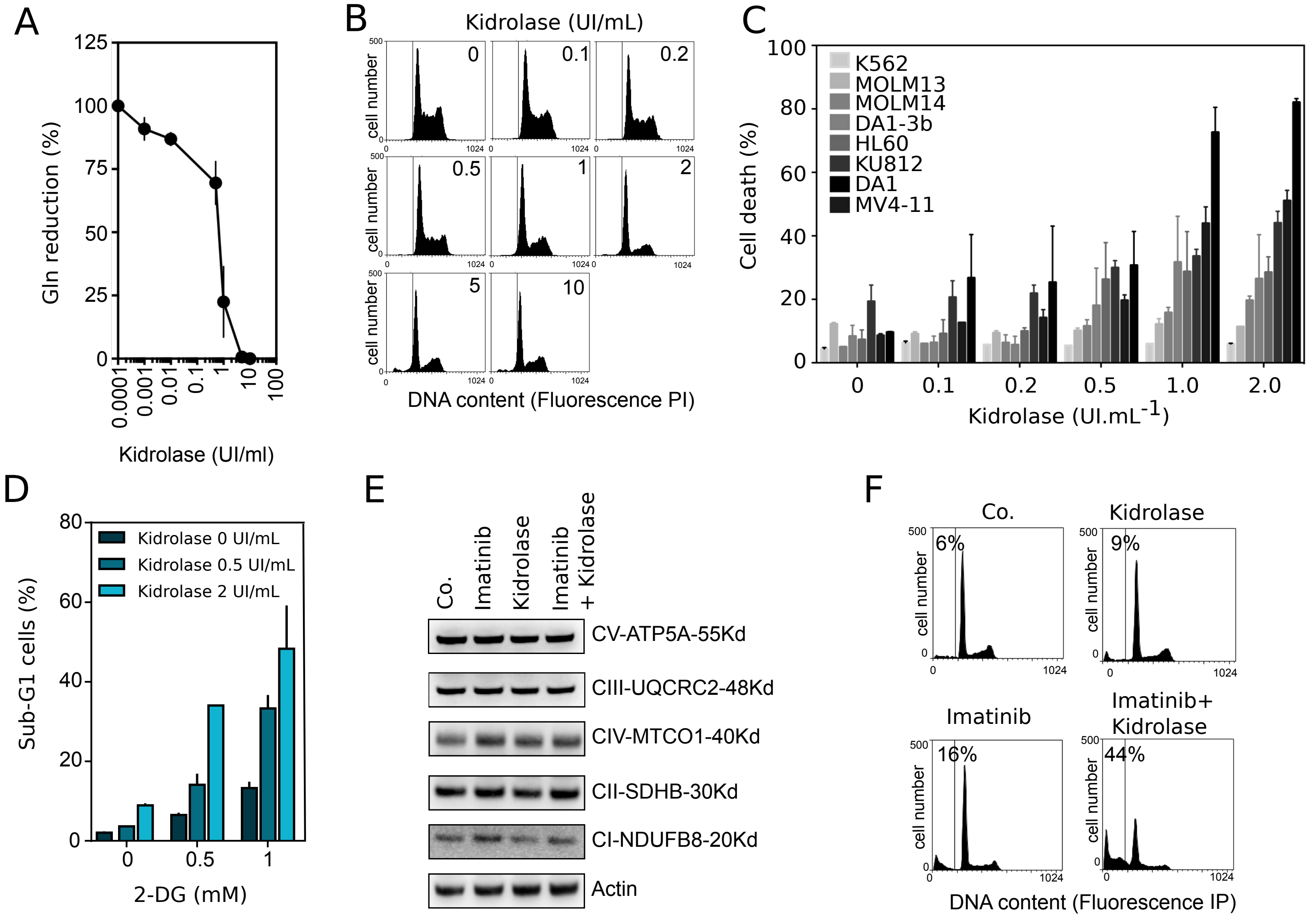


supplementary Figure 3
